# Dissecting the Genetic Basis of Superior Traits in Thermosensitive Genic Male Sterile Line 1892S Through Genome-Wide Analysis

**DOI:** 10.1101/2024.07.29.605629

**Authors:** Wei Zhang, Dewen Zhang

## Abstract

Hybrid rice has revolutionized food security by leveraging heterosis, a phenomenon where offspring outperform their parents. Sterile lines, crucial for controlled cross-pollination in hybrid breeding, have played a central role in this success. This study delves into the superior alleles of 1892S, a two-line sterile rice line widely used as a female parent in central China. By integrating extensive hybridization data, high-throughput genome sequencing, and bioinformatic analysis (RiceNavi), we elucidate the genetic underpinnings of 1892S’s exceptional adaptability. The results reveal its remarkable compatibility as a female parent in over 114 hybrid varieties, potentially due to the influence of Japonica characteristics contributing to strong hybrid vigor. Furthermore, some favorable alleles were identified that were associated with lodging resistance, high yield potential, and improved nitrogen use efficiency etc. The comprehensive characterization of 1892S provides valuable insights for future hybrid rice breeding programs, ultimately facilitating the development of superior rice varieties.

## Introduction

Heterosis, or hybrid vigor, plays a critical role in boosting crop yield and quality(Paril et al. 2024; Wu et al. 2021). The discovery and successful application of thermosensitive genic male sterile lines (TGMS Lines) in rice revolutionized hybrid rice breeding for self-pollinating crops (Wang et al. 2005; Yuan 2014; Ashraf et al. 2020). TGMS lines like 1892S offer several advantages over traditional sterile lines used in three-line breeding systems (Yang Lian-song and Yi-song 2012; Lian-song et al. 2016). These advantages include wider cross-compatibility, simpler sterile material reproduction, higher hybrid seed production yield, and easier sterility gene transfer (Lian-song et al. 2016; Yang Lian-song and Yi-song 2012).

The risk and slow rate of popularization of two-line hybrid rice are mainly due to the unstable sterility and poor combining ability of some indica sterile lines, and the limited production and application of the F_1_ hybrid (Xu et al. 2023; Cao and Zh 2014). However, the two-line sterile line 1892S stands out for its exceptional characteristics. It exhibits high combining ability, good propagation traits, strong resistance to diseases, and excellent grain quality. Additionally, 1892S-based hybrids demonstrate strong heterosis, tolerance to fertilizers, resistance to lodging, and broad adaptability(Khan et al. 2024; Lian-song et al. 2016).

Despite its success, a comprehensive analysis of 1892S’s genetic makeup is lacking. This knowledge gap hinders the wider application of 1892S and limits its potential to optimize breeding programs in the era of molecular design breeding, which relies heavily on in-depth varietal analysis for the efficient development of new lines (Collard et al. 2008).

This study aims to address this critical gap by employing a multi-pronged approach. We will leverage RiceNavi, a powerful rice breeding tool that provides detailed quantitative trait gene information, to analyze the excellent alleles present within the entire genome of 1892S (Wei et al. 2021; Qianlong et al. 2023). Additionally, we will assemble the genome of 1892S to support the existence of these beneficial genes. Furthermore, we will analyze the Indica-Japonica properties of 1892S using indel markers to gain insights into its heterosis characteristics(Shen et al. 2004).

The comprehensive analysis empowers breeders with a detailed genetic analysis of 1892S. it reveals favorable alleles associated with crucial agronomic traits like lodging resistance, high yield potential, and improved nitrogen use efficiency. Armed with this knowledge, breeders could make strategic decisions about crossing partners, breeding process optimization, and prioritizing desired traits in the development of hybrid rice varieties.

## Results

### Phenotypic Characterization of 1892S

1892S exhibited several notable phenotypic characteristics upon cultivation. Such as 1892S displayed visibly stronger stems compared to other varieties. Its hybrid variety has potentially higher yields with increased grain number per spike. Its high stigma exsertion rate which could improves pollination efficiency by ensuring better exposure of the stigma to pollen grains (Y.N.J. et al. 2022). Additionally, as an indica-type variety, 1892S demonstrates compatibility with other indica varieties, facilitating hybridization to leverage complementary traits and achieve superior agronomic advantages. This highlights 1892S’s potential as a valuable genetic resource for breeding programs focused on developing high-yielding and resilient rice varieties suited to specific environmental conditions.

### Functional Locus Analysis of 1892S

High-throughput sequencing of 1892S was performed, achieving comprehensive coverage of the genome (110x) and an impressive Q30 value exceeding 92.69%. Clean data was used for further analysis with the rice reference genome Nipponbare (MSU 7.0) as a reference. RiceNavi software was employed to identify quantitative trait nucleotides (QTNs) associated with various phenotypic traits (Wei et al. 2021). This analysis identified a total of 319 QTN loci across the 1892S genome. The predicted effects of these QTNs on the phenotype of 1892S were then interpreted (Table S1). This comprehensive analysis revealed 32 potentially superior genes within 1892S (Table 1). This information provides valuable insights into the genetic basis of 1892S’s traits and paves the way for further investigation and breeding efforts to improve rice varieties.

### Genome Assembly and Assessment

The assembled genome of 1892S has an estimated size of 324.7 Mb with a GC content of 43.00%. The longest contig fragment is 147,639 bp, while the Contig N50 and N90 values are 16,196 bp and 3,277 bp, respectively (Table 2). The completeness of the assembly was assessed using BUSCO, revealing that 93.8% of the core genes were successfully identified, with only 2% missing (Table 3). Synteny analysis confirmed a high degree of similarity between the 1892S genome and the reference genome MSU 7.0. Overall, the 1892S genome assembly demonstrates high reliability, completeness, and synteny with the reference, providing a solid foundation for further studies (Figure 1).

**Table 2.** Preliminary genome assembly of 1892S.

| Type | Contigs |
| --- | --- |
| Total length (bp) | 324669546 |
| GC (%) | 43.00 |
| N50 (bp) | 16196 |
| N90 (bp) | 3277 |
| Longest (bp) | 147639 |
| contigs | 45720 |
| contigs ($\geq 10000$ bp) | 10273 |
| contigs ($\geq 25000$ bp) | 2671 |
| contigs ($\geq 50000$ bp) | 375 |

**Table 3.**
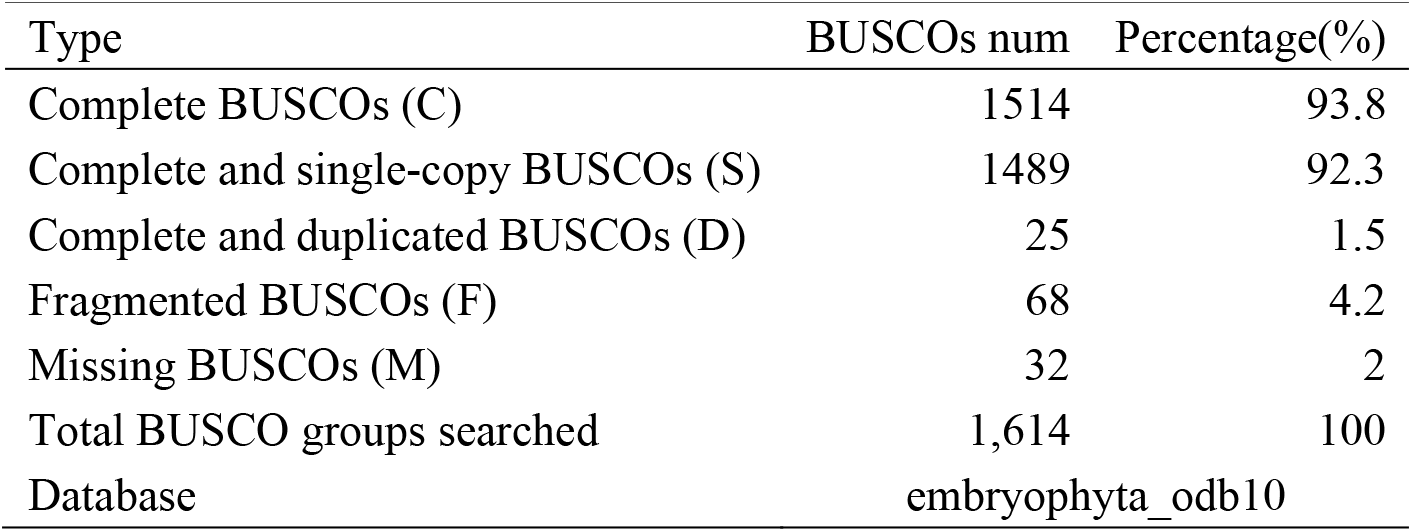
Quantitatively assessed of 1892S genome assembly integrity.

| Type | BUSCOs num | Percentage(%) |
| --- | --- | --- |
| Complete BUSCOs (C) | 1514 | 93.8 |
| Complete and single-copy BUSCOs (S) | 1489 | 92.3 |
| Complete and duplicated BUSCOs (D) | 25 | 1.5 |
| Fragmented BUSCOs (F) | 68 | 4.2 |
| Missing BUSCOs (M) | 32 | 2 |
| Total BUSCO groups searched | 1,614 | 100 |
| Database | embryophyta_odb10 |  |

**Figure 1.**
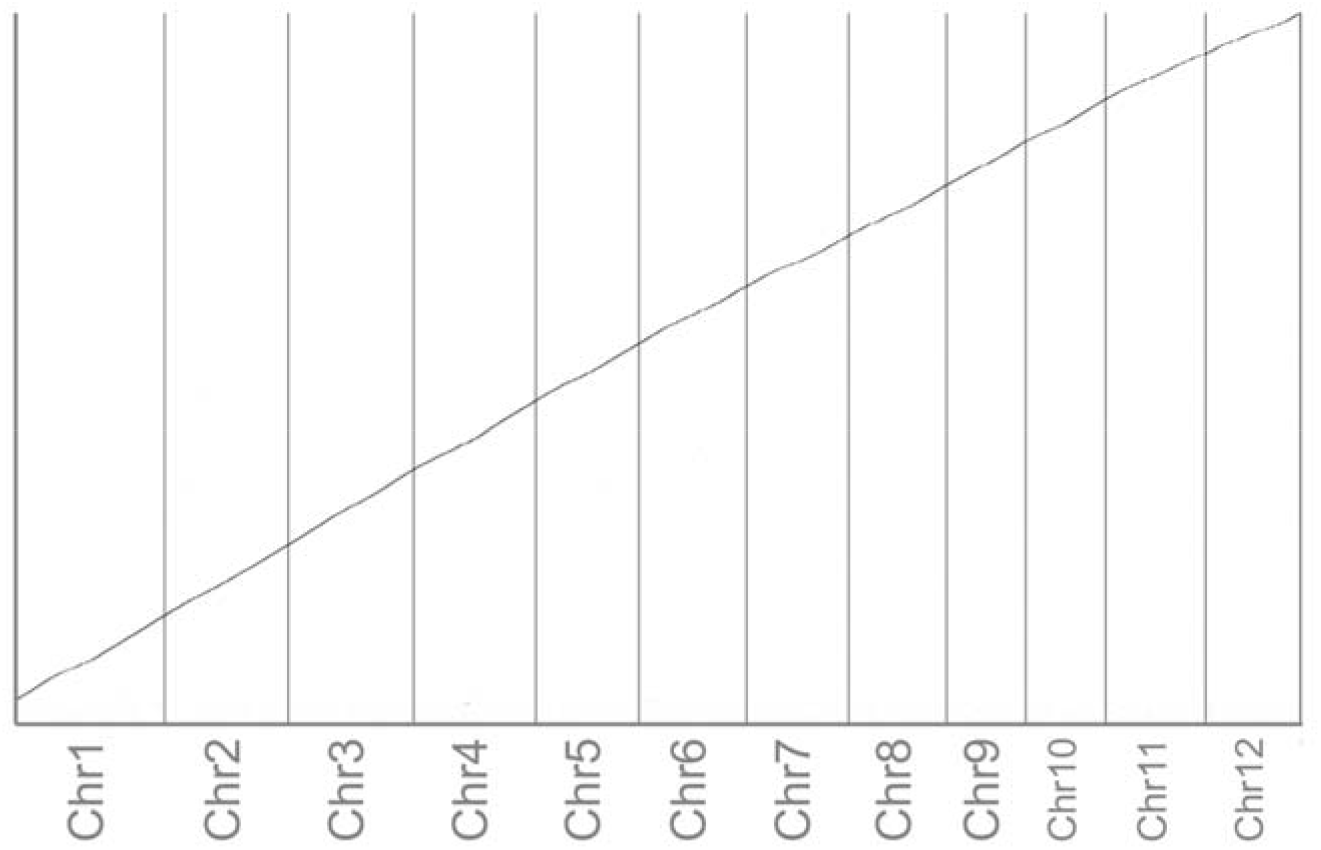
Sequence collinearity analysis among MSU 7.0 and 1892S

### Local Alignment of Gene Sequences

To determine the location of contigs within the reference genome and gain context about the assembled sequences, the draft genome of 1892S was mapped to the reference genome using Minimap2. Extracted gene sequences and their corresponding contig sequences from the draft assembly were then aligned and compared using MUSCLE software. Based on the alignment results, genes within the 1892S draft genome exhibiting high similarity with known superior alleles were identified as excellent genes (Table 1). This successful comparison confirms the presence of these excellent genes within the draft genome of 1892S.

### Indica-Japonica Classification Using InDel Markers

Indel markers were employed to distinguish between the Indica and Japonica subspecies of rice and understand the genetic relationship between 1892S and these varieties. Briefly, BAC clone sequences were aligned with the genome of rice variety *93-11* to identify matching regions. Primer sequences flanking indel markers were designed to amplify and sequence these regions. A total of 43 indel markers specific to the BAC clone were identified and mapped to the *93-11* genome. The 1892S contig sequence was then aligned to the *93-11* genome, and corresponding sequences were extracted based on the marker positions. Analysis of these alignments revealed 30 indel markers present in all three varieties (Table 4). Six of these markers indicated shared sequences between 1892S and Nipponbare (*Japonica*), while the remaining 24 matched *93-11* (Indica). These findings suggest that 1892S possesses a mixed genetic background with characteristics of both Indica and Japonica subspecies.

**Table 4.**
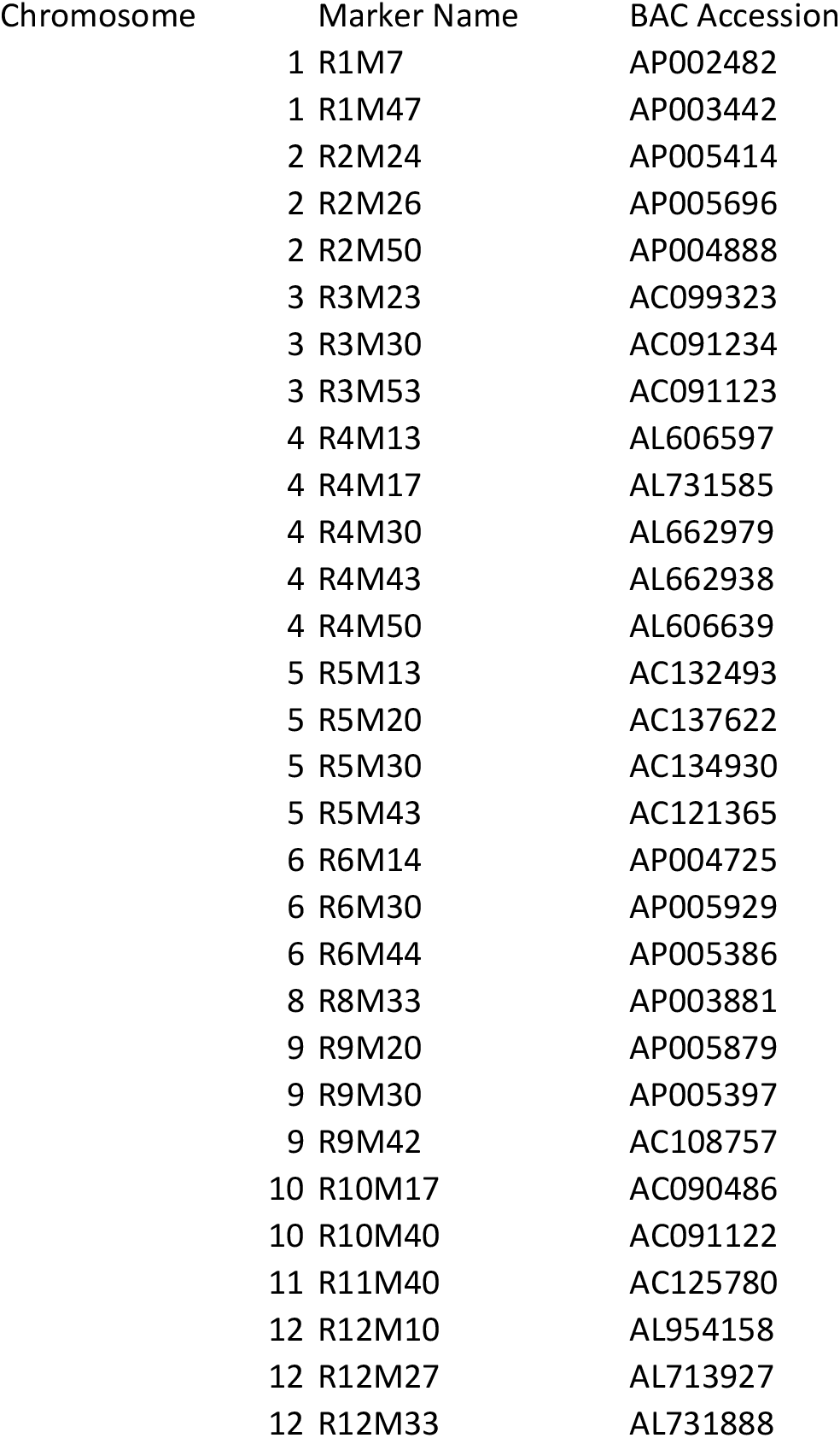
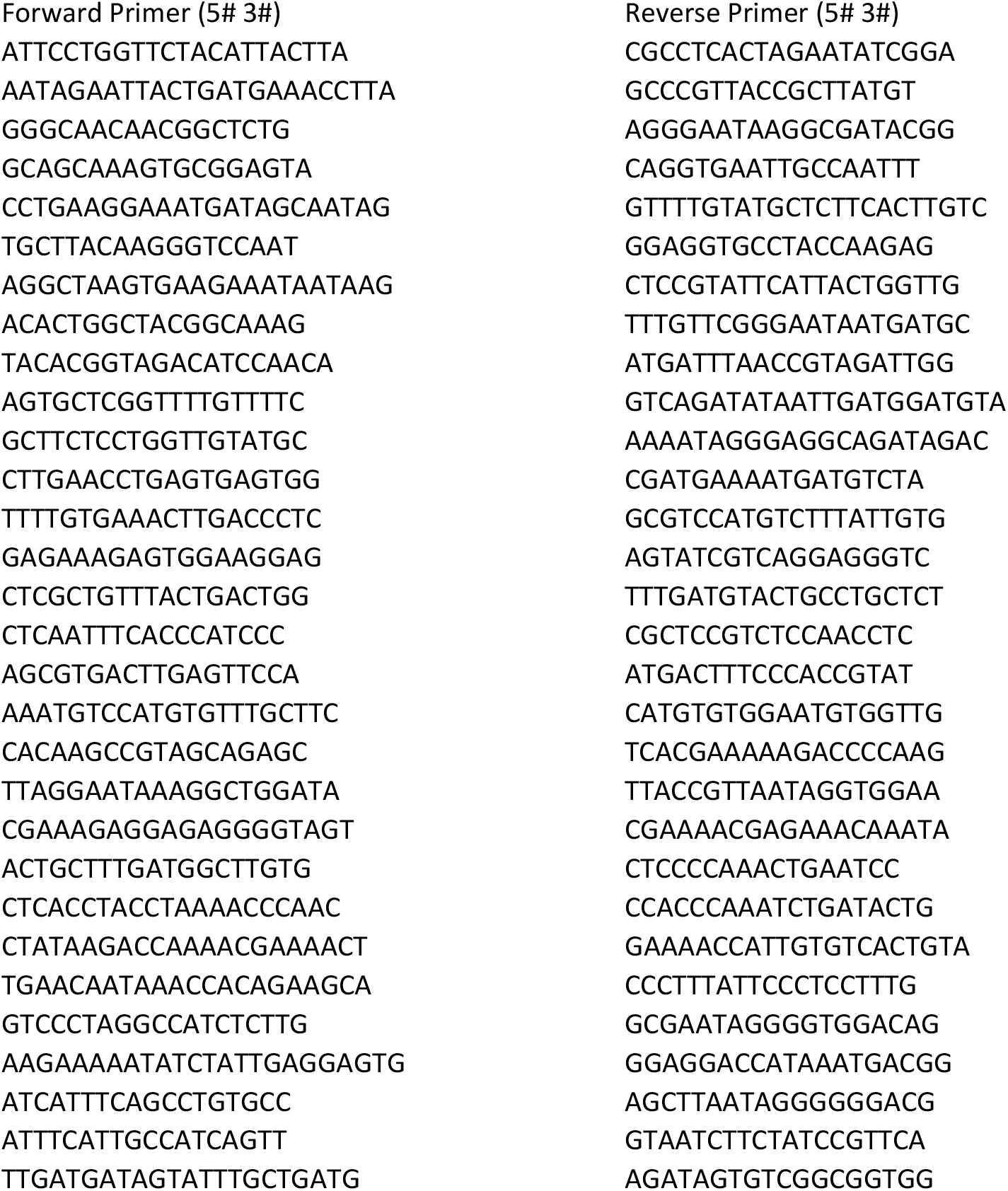
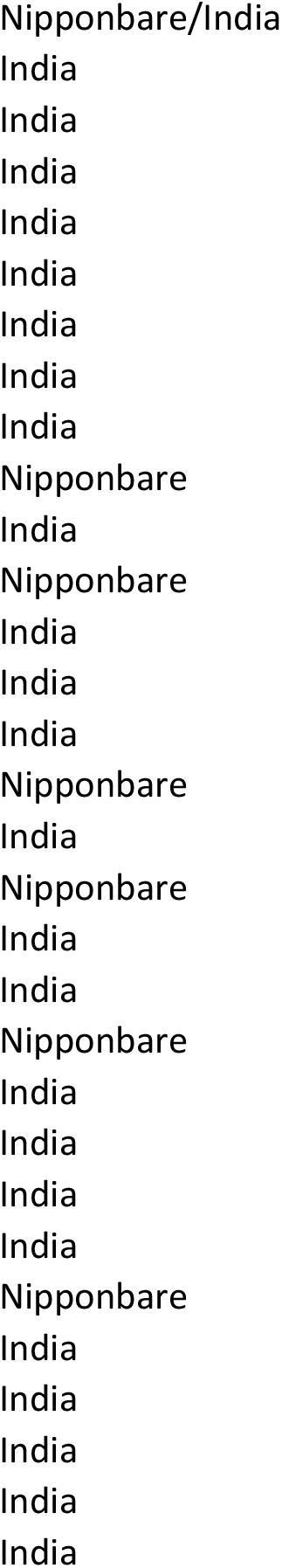
The comprehensive information on the InDel markers designed for diverse combinations of japonica and indica.

### Favorable Traits and Breeding Potential of 1892S

This section combines findings from several subsections to highlight the key advantages and breeding potential of the 1892S rice variety.

Wide Affinity: The wide affinity *S5* locus in 1892S, as indicated in Table 1, aligns with previous studies highlighting its advantageous traits. Specifically, the *S5-n* gene plays a crucial role in controlling the wide affinity of rice intersubspecific hybrids. Loss of sequence in the *S5-n* gene results in loss of its function, yet hybrid-wide affinity remains unaffected with both indica and japonica rice varieties (Chen et al. 2008). To confirm the presence of the *S5-n* gene in 1892S, the assembled contig sequence of its genome was locally compared with the corresponding site in the reference genome. The comparison revealed a deletion in the N-terminal region, providing evidence that 1892S indeed possesses the *S5-n* gene (附件 S5.clw).

The extensive use of 1892S, with its wide affinity gene *S5-n*, is evident in the approval of combination varieties where it serves as the female parent. A total of 114 hybrid rice varieties with 1892S as the female parent have received approval. Notably, since its technical appraisal in Anhui Province in 2004, the first variety, “Wandao 153,” incorporating 1892S as the female parent, obtained approval the following year, marking a significant milestone in its contribution to national rice variety selection (Liansong and Yisong 2006). Over the years, from 2005 to 2023, there has been a consistent increase in the approval of hybrid rice varieties with 1892S as the female parent, culminating in a record-setting 24 approvals in 2021. These varieties are widespread and distributed across 9 provinces, with 42 provincial rice varieties receiving national approval. Among these, the number of varieties approved in Anhui ranks second only to those with national approval. This analysis underscores the broad affinity and significant contribution of 1892S to rice variety development.

Lodging Resistance: 1892S possesses the APO1 site linked to lodging resistance (Table 1). This finding aligns with observations from approved hybrid combinations using 1892S, which exhibit anti-inversion characteristics. Additionally, the presence of semi-dwarf trait genes (SBI/Sd1/Ghd7 and SD1/OsSPL14) in 1892S contributes to its compact growth habit (Liu et al. 2018; Asano et al. 2007; Jiao et al. 2010), a factor not only enhancing lodging resistance but also allowing for higher planting density and potentially greater yield. Real-world examples like Wandao 153 further solidify 1892S’s contribution to lodging resistance. This variety demonstrates exceptional resilience, characterized by short stature, robust stems, and superior root architecture (Miaomiao et al. 2013).

High Yield Potential: The genetic makeup of 1892S includes genes associated with a high number of grains per panicle, including *Gn1a/OsCKX2* (Ashikari et al. 2005), *GNP1/OsGA20ox1* (Wu et al. 2016), and *An-2/OsLOGL6/LABA1* (Table 1) (Gu et al. 2015; Hua et al. 2015). These genes play a crucial role in regulating grain number and panicle development, ultimately impacting yield per unit area. Varieties derived from 1892S exhibit a significantly higher number of grains per spike and demonstrably higher yields compared to controls.

Improved Nitrogen Use Efficiency: The presence of *OsNR2* and *NRT1.1B* sites in 1892S suggests enhanced potential for nitrogen use efficiency (NUE) (Table 1) (Gao et al. 2019; Hu et al. 2015). The indica *OsNR2* variant offers superior traits compared to its japonica counterpart, leading to increased chlorate sensitivity, improved nitrate uptake, and ultimately, greater grain yield (Gao et al. 2019). *NRT1.1B* also plays a significant role in nitrate signaling and NUE. By harboring these genes, 1892S holds promise for optimizing nitrogen utilization in rice crops.

Sterile Line Development: 1892S serves as a valuable resource for breeding two-line sterile rice varieties. As a two-line sterile line itself, 1892S has been used to develop many additional sterile lines. Notably, 1892S exhibits a high stigma exposure rate, a characteristic crucial for efficient hybridization (Y.N.J. et al. 2022). Gene sequence analysis confirmed the presence of stigma exsertion genes (gs3/gw8/gs9) within 1892S, further supporting its role in sterile line development (Zhu et al. 2023).

## Discussion

The success of molecular design breeding relies heavily on the accurate understanding of the genetic makeup of the crop, the traits of interest, and the complex interactions between genes and the environment (Ahmar et al. 2020; Begna). It can significantly enhance the efficiency and effectiveness of crop improvement programs (Pasala et al. 2024). This approach has significant advantages in crop breeding, where molecular design breeding allows breeders to work at the genetic level to select and combine genes with specific traits very precisely to create new varieties with the desired traits (Jeon et al. 2023). Before proceeding with molecular breeding, it is essential to understand the characteristics of the variety (Moose and Mumm 2008). This includes an in-depth understanding of the genetic background, agronomic traits, growth habits, stress tolerance, quality characteristics of the target crop, etc. This information can help determine breeding goals, select suitable parents, and predict trait performance in crossbred offspring. The success of molecular breeding depends to a large extent on the accurate understanding and utilization of the characteristics of the variety (Ahmar et al. 2020).

As a rice variety widely used in actual production, the association between the characteristics of 1892S and specific functional gene loci may not be fully understood. Although modern molecular breeding techniques have come a long way, there are still some challenges (Lamichhane and Thapa 2022). To better understand the association between the characteristic properties of 1892S and specific functional gene loci, this study analyzed the *Indica*-*Japonica* properties of 1892S using an improved indel maker analysis strategy. Due to the distant genetic relationship between indica rice and japonica rice, indica-japonica rice hybridization often produces many genetic recombination and segregation types, which provides breeders with more selection opportunities (Xu et al. 2020). The results showed that 1892S had a certain proportion of japonica rice attributes, and the characteristics of 1892S, as the female parent had strong stress resistance, tillering ability, and yield advantages, confirmed the results of this analysis (Table 3).

This study implemented an enhanced RiceNavi approach. First, RiceNavi was used to predict the characteristic features of 1892S. Second, the assembled 1892S genome sequence (second-generation) was used to confirm the presence of gene sequences predicted by RiceNavi. This approach enhances the accuracy of variety characterization and analysis. For example, analyzing the wide affinity of 1892S solely through sequence alignment might only suggest the presence of the S5-n gene due to a deletion in the N-terminal region. However, our analysis not only predicted a wide affinity gene in 1892S but also confirmed this through real-world production data. This is supported by the fact that 114 varieties utilizing 1892S as the female parent have been nationally approved across nine provinces. These varieties have diverse paternal parents originating from different ecological regions and genetic backgrounds, further supporting the analysis of wide affinity genes in 1892S.

Furthermore, this study predicted lodging resistance in 1892S, and analysis confirmed the presence of loci related to this trait, such as APO1 site SCM2 (Ookawa et al. 2010), SBI/, and SD1 (Guha et al. 2024). Hybrid varieties with 1892S as the female parent also demonstrate strong lodging resistance, exemplified by Wandao 153 – a widely grown variety in the middle and lower Yangtze River region known for its exceptional lodging resistance.

While next-generation sequencing technologies offer high-resolution data for genome-wide analysis, limitations exist, including challenges in covering specific genomic regions and detecting large structural variants or repeats (Satam et al. 2023). Advancements in sequencing technology, particularly PacBio SMRT and Oxford Nanopore platforms, offer longer read lengths and higher accuracy, facilitating a more comprehensive analysis of genomic structures, including complex regions and variations in repeats (Udaondo et al. 2021). Utilizing data from multiple generations alongside novel analytical methods will allow for more precise identification and verification of functional gene loci in 1892S, providing deeper support for molecular design breeding. This data can empower breeders to gain a clearer understanding of 1892S’s genetic background, refine cross-breeding strategies, and enhance the efficiency and success rate of breeding new varieties. Additionally, integrating multi-omics data offers a more comprehensive view of gene regulation across different levels, leading to more precise guidance for molecular design breeding (Zhang et al. 2022). As technology advances and costs decline, next-generation sequencing and other multi-omics technologies will play an increasingly significant role in propelling molecular design breeding forward (Mahmood et al. 2022; Yang et al. 2021).

Overall, this study conducted a comprehensive analysis of 1892S’s characteristic features, revealing its dominance in heterosis, wide affinity, lodging resistance, and other traits. Additionally, we identified some potentially disadvantaged genes in 1892S. This comprehensive characterization not only provides theoretical support for the wider adoption and application of 1892S but also establishes a solid foundation for using 1892S as breeding or research material in future endeavors.

## Material and methods

### Rice materials

The Rice Research Institute of the Anhui Academy of Agricultural Sciences bred the TGMS line 1892S. The passing of the technical appraisal in 2004 and the application for new plant variety rights in the same year, followed by authorization in 2007, the variety right number is CNA20040612.4 (https://www.ricedata.cn).

### DNA Extraction and Genome Sequencing

The genomic DNA extraction from the young leaves of 1892S plants using the CTAB method followed by sequencing library construction. Utilizing the HiSeq2500 next-generation sequencing platform with a sequencing depth of 100 ensures thorough coverage of the genome. The generation of 157,494,513 RawReads reflects the extensive data obtained from the sequencing process, enabling in-depth genome-wide analysis and identification of key genetic features and variations associated with 1892S.

### Functional locus analysis using RiceNavi

Use Trimmomatic to trim and filter the raw reads based on quality scores and adapter sequences. Then run FastQC on the trimmed read files to assess their quality and identify any remaining issues or biases. These software bowtie2, samtools, sambamba, GATK3, GATK4, Manta, bam2fastq are called by RiceNavi (Wei et al. 2021). The rice genome (MSU v7) was downloaded from http://rice.uga.edu/. The whole genome clean data was Aligned to the reference genome. Utilize RiceNavi-QTNpick mode to calculate genotyping causal variant sites based on the alignment results. The mode provides information on QTN sites and their associated characteristics based on rice accessions in QTNlib with different alleles of each causative site. The detailed information on the QTN site comes from the data from Wei’s article (Wei et al. 2021).

### Genome assembly and local alignment of gene sequences

The genomic clean data were used to assemble the genome by SOAPdenovo2 (Luo et al. 2012). Utilize GapCloser to fill the gaps in the assembled genome using the paired relationships of short reads. Run BUSCO on the assembled genome to assess its integrity and completeness by comparing it to conserved single-copy orthologs (Manni et al. 2021). Perform collinearity analysis between the assembled genome and the reference genome using Mummer software which was used to identify large-scale structural variations and similarities between the assembled genome and the reference genome (Marcais et al. 2018). Align the assembled genome to the reference genome using Minimap2 (Li 2021). The gene sequence of the reference genome is downloaded from http://rice.uga.edu/. Muscle was used to align the gene sequences locally and assembled contig sequences to identify similarities, differences, and structural variations at the gene level (Edgar 2022).

### Indica-japonica attribute identification

Some Bacterial Artificial Chromosome (BAC) sequences that come from Shen’s article (Shen et al. 2004) were downloaded from the NCBI database. These sequences are likely to contain the InDel markers of interest. The downloaded BAC sequences were aligned to the genome of rice variety 93-11 using the Minimap2. Based on the mapping results, the corresponding sequences were extracted. These sequences should contain the InDel markers. Forward and Reverse Primers specific to the InDel markers were used to extract the sequences from both indica and japonica rice varieties separately. The extracted InDel sequences from India and japonica were mapped back to the 93-11 genome to provide detailed information about their positions and contexts within the genome. Contigs of 1892S were also mapped to the 93-11 genome using Minimap2, which was used to find corresponding regions in 1892S that align with the InDel markers identified in 93-11. By combining the mapping information from the previous steps, the sequences of the InDel markers from 1892S were obtained. These sequences were used for comparison with those from indica and japonica. The sequences of InDel markers from indica, japonica, and 1892S were aligned together using MUSCLE. it is determined whether each sequence of InDel marker from 1892S is more like the indica or japonica variety.

### Data analysis of hybrid rice combinations with 1892S

To analyze the hybrid rice combinations involving 1892S as the female parent, we visited the National Rice Data Center website (https://www.ricedata.cn/) to access the data on hybrid rice combinations for searching for hybrid rice varieties which 1892S was listed as the female parent. The relevant information was extracted, such as the names, approval numbers, and characteristics of the hybrid rice varieties. We also Gathered information on TGMS lines that 1892S was the female parent.

## Conclusion

In conclusion, the 1892S demonstrates a unique combination of traits favorable for breeding high-yielding, adaptable, and resource-efficient rice varieties. Its wide affinity, lodging resistance, high yield potential, improved NUE, and role in sterile line development make it a valuable resource for future rice breeding programs.

